# SUMOylation of the m6A reader YTHDF2 by PIAS1 promotes viral RNA decay to restrict EBV replication

**DOI:** 10.1101/2023.08.08.552509

**Authors:** Febri Gunawan Sugiokto, Farjana Saiada, Kun Zhang, Renfeng Li

## Abstract

YTHDF2 is a member of the YTH protein family that binds to N6-methyladenosine (m6A)-modified RNA, regulating RNA stability and restricting viral replication, including Epstein-Barr virus (EBV). PIAS1 is an E3 SUMO ligase known as an EBV restriction factor, but its role in YTHDF2 SUMOylation remains unclear. In this study, we investigated the functional regulation of YTHDF2 by PIAS1. We found that PIAS1 promotes the SUMOylation of YTHDF2 at three specific lysine residues (K281, K571, and K572). Importantly, PIAS1 enhances the antiviral activity of YTHDF2, and SUMOylation-deficient YTHDF2 shows reduced anti-EBV activity. Mechanistically, YTHDF2 lacking SUMOylation exhibits reduced binding to EBV transcripts, leading to increased viral mRNA stability. Furthermore, PIAS1 mediates SUMOylation of YTHDF2’s paralogs, YTHDF1 and YTHDF3. These results collectively uncover a unique mechanism whereby YTHDF2 controls EBV replication through PIAS1-mediated SUMOylation, highlighting the significance of SUMOylation in regulating viral mRNA stability and EBV replication.

**Importance:** N6-methyladenosine (m6A) RNA modification pathway plays important roles in diverse cellular processes and viral life cycle. Here, we investigated the relationship between PIAS1 and the m6A reader protein YTHDF2, which is involved in regulating RNA stability by binding to m6A-modified RNA. We found that both the N-terminal and C-terminal regions of YTHDF2 interact with PIAS1. We showed that PIAS1 promotes the SUMOylation of YTHDF2 at three specific lysine residues. We also demonstrated that PIAS1 enhances the anti-Epstein-Barr virus (EBV) activity of YTHDF2. We further revealed that PIAS1 mediates the SUMOylation of other YTHDF family members, namely YTHDF1 and YTHDF3, due to their similarities with YTHDF2. These findings together illuminate an important regulatory mechanism of YTHDF2 in controlling viral RNA decay and EBV replication through PIAS1-mediated SUMOylation.

## Introduction

Host restriction factors play a crucial role in defending against viral infection, replication, and egress by targeting viral proteins, viral DNA, and/or RNA (1). One restriction factor YTH N6-methyladenosine RNA binding protein 2 (YTHDF2) is an RNA-binding protein that specifically recognizes and binds to N6-methyladenosine (m6A)-modified RNA (2, 3). Studies from our lab and other groups have highlighted the key role of YTHDF2 in limiting the reactivation of Epstein-Barr virus (EBV) and Kaposi’s sarcoma-associated herpesvirus (KSHV) through targeting and destabilizing viral and cellular RNAs (4-7). YTHDF2 has also been implicated in destabilizing hepatitis B virus mRNA in coordination with ISG20 (8). Our recent study showed that caspase-mediated cleavage of YTHDF2 antagonize its anti-viral activity during EBV reactivation process (7).

In addition to cleavage, YTHDF2 can also be regulated through phosphorylation, ubiquitination, and SUMOylation. Phosphorylation of YTHDF2 at S39 and T381 by the EGFR/SRC/ERK signaling pathway contributes to its stabilization (9). The stability of YTHDF2 is also regulated by the activity of cyclin-dependent kinase 1 (CDK1), where phosphorylated YTHDF2 can be targeted for ubiquitination-dependent degradation by E3 ubiquitin ligase complexes, including Cullin 1 (CUL1), Cullin 4A (CUL4A), damaged DNA-binding protein 1 (DDB1), and S-phase kinase-associated protein 2 (SKP2) (10). These studies highlight the significance of post-translational modifications in modulating the function of YTHDF2 in mRNA regulation, particularly in the context of m6A modification.

SUMOylation is a post-translational modification (PTM) process involving the attachment of small ubiquitin-like modifier (SUMO) proteins (SUMO1/2/3) to target proteins. This process relies on a cascade of enzymes, including the E1 activating enzyme complex SAE1/SAE2, the E2 conjugating enzyme UBC9, and specific E3 protein ligases (11). Among the E3 ligases, the Protein Inhibitor of Activated family proteins (PIAS1, PIASx/PIAS2, PIAS3, PIASy/PIAS4) are known to be involved in SUMOylation (12).

Our previous studies have identified PIAS1 as a restriction factor for EBV (13) and as an E3 ligase to enhance the anti-EBV activity of SAMHD1 through SUMOylation (14). In this study, we demonstrated that PIAS1 interacts with YTHDF2 to promote the anti-EBV activity of YTHDF2. We showed that PIAS1 promotes YTHDF2 SUMOylation at three lysine residues, K281, K571, and K572. The interaction between YTHDF2 and PIAS1, as well as the SUMOylation process mediated by PIAS1, is extended to its paralogs YTHDF1 and YTHDF3, highlighting the significance of PIAS1-mediated SUMOylation in the regulation of m6A readers and the m6A RNA modification pathway.

## Results

### PIAS1 promotes YTHDF2 SUMOylation

In our previous studies, we have shown that both YTHDF2 and PIAS1 play crucial roles in restricting EBV replication (7, 13, 15). YTHDF2 contributes to the decay of viral and cellular genes, while PIAS1 inhibits viral gene transcription. PIAS1, as an E3 SUMO ligase, is responsible for the SUMOylation of various proteins. Recently, we have discovered that PIAS1 promotes the SUMOylation of SAMHD1 on multiple lysine residues to enhance its anti-viral activity (14).

A recent study identified YTHDF2 as a SUMOylated protein (16). However, the E3 ligase responsible for YTHDF2 SUMOylation is not known. We hypothesize that PIAS1 functions a an E3 ligase facilitating YTHDF2 SUMOylation. To test our hypothesis, we transfected HEK293T cells with plasmids expressing YTHDF2, PIAS1 and SUMO2. We found that transfection of SUMO2 or PIAS1 slightly increases the total SUMOylation level (**Figure 1A**, top panel, lanes 2 and 3 vs 1) while SUMO2 and PIAS1 co-transfection significantly promote SUMOylation (**Figure 1A**, top panel, lane 4). To determine whether PIAS1 promotes th SUMOylation of YTHDF2, we then immunoprecipitated (IPed) YTHDF2 by anti-V5 antibody-conjugated magnetic beads. Western blot analysis with anti-SUMO2/3 antibody showed that YTHDF2 SUMOylation is increased when SUMO2 or PIAS1, or SUMO2 and PIAS1 are expressed (**Figure 1B**, top panel, lanes 2-4 vs 1), suggesting YTHDF2 is targeted for SUMOylation by PIAS1.

**Figure 1.**
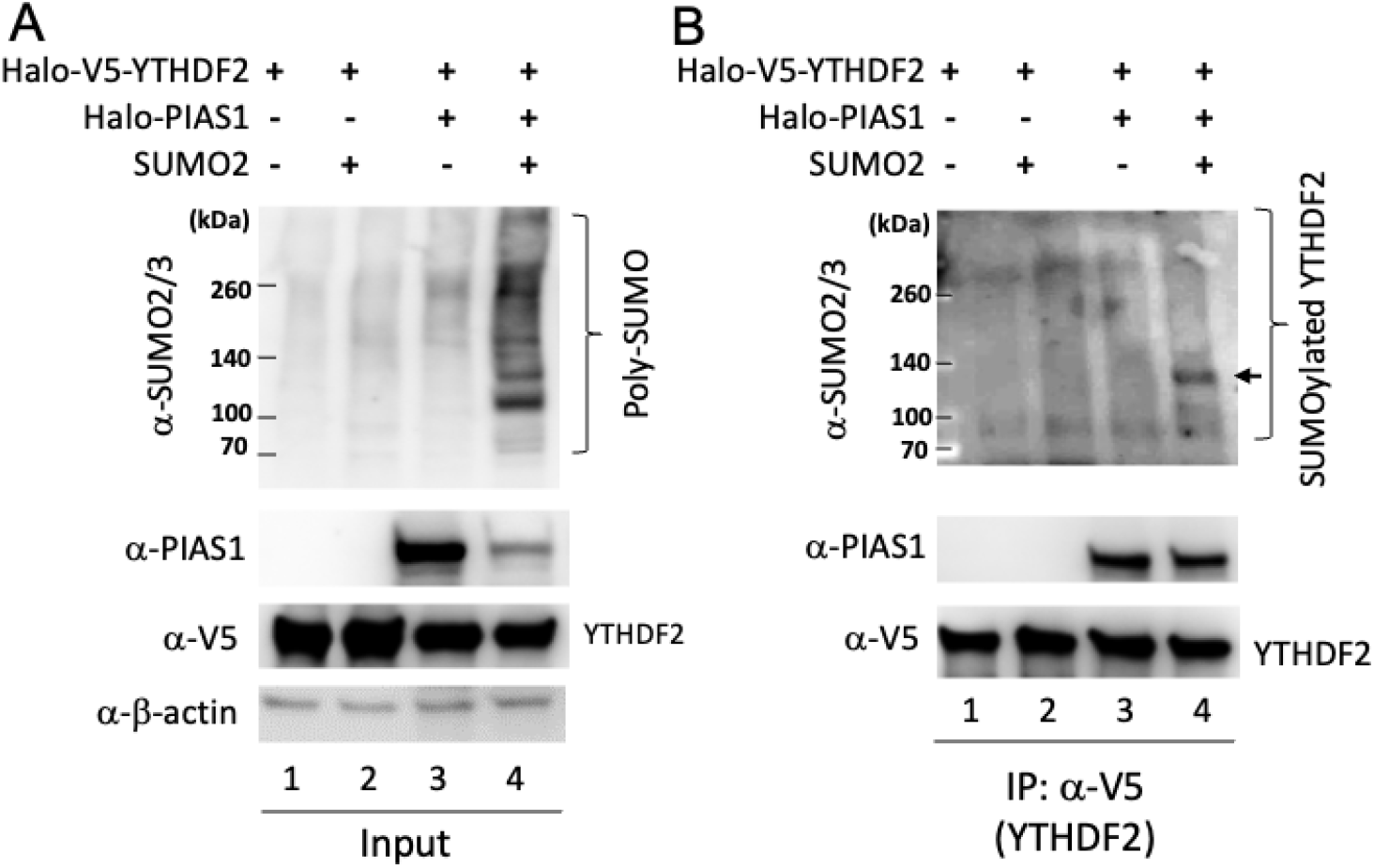
PIAS1 promotes YTHDF2 SUMOylation. HEK293T cells were transfected with Halo-V5-YTHDF2, Halo-PIAS1 and SUMO2 plasmids as indicated. **A.** Western Blot (WB) were performed for whole cell lysate (Input) using anti-SUMO2/3, anti-PIAS1, anti-V5, and anti-β-actin antibodies. Bracket denotes SUMOylated proteins. **B.** YTHDF2 was immunoprecipitated with anti-V5 magnetic beads and WB was performed using antibodies as indicated. **C.** Bracket denotes SUMOylated YTHDF2.

### PIAS1 interacts with YTHDF2

When YTHDF2 is IPed by anti-V5 beads, we noticed that PIAS1 is also co-immunoprecipitated (Co-IPed) with the beads (**Figure 1B**, PIAS1 blot, lanes 3 and 4), indicating an interaction between PIAS1 and YTHDF2. To further determine whether YTHDF2 interacts with PIAS1 and which regions within YTHDF2 are responsible for this interaction, we transfected HEK293T cells with plasmids expressing full-length PIAS1 and full-length or individual fragments of YTHDF2 (**Figure 2**). We then performed Co-IP experiments and found that PIAS1 is strongly Co-IPed by full-length YTHDF2 (**Figure 2B**, top panel, lane 1). The Co-IP of PIAS1 was also observed in all YTHDF2 fragments except the central region of YTHDF2 (aa 167-367) (**Figure 2B**, top panel, lanes 2, 3, 5 and 6 vs lane 4). These results suggested that both N-terminal (aa 1-166) and C-terminal (aa 368-579) of YTHDF2 bind to PIAS1.

**Figure 2.**
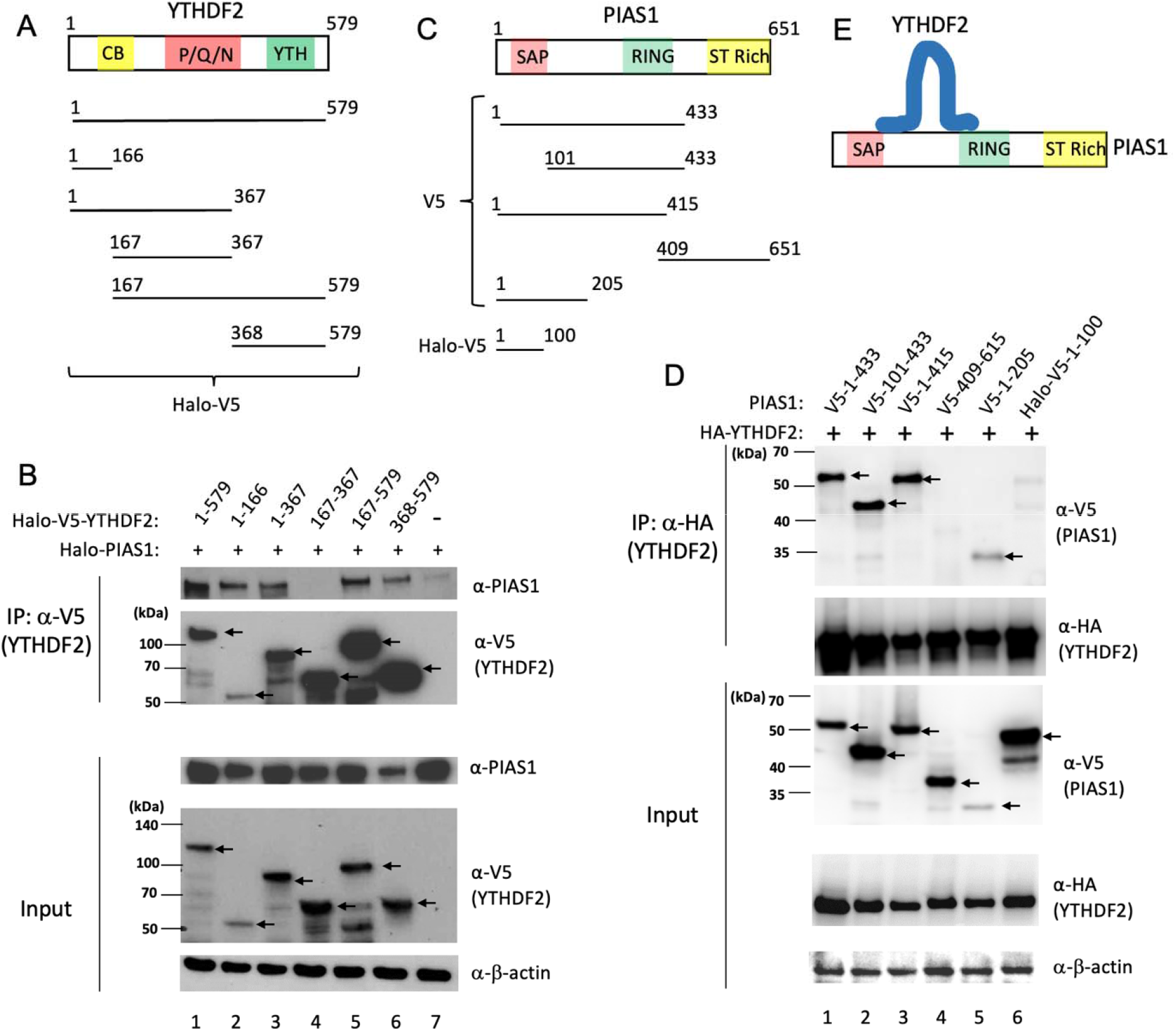
PIAS1 interacts with YTHDF2. Related to Figure S1. **A.** The illustration of full length YTHDF2 (1-579) or YTHDF2 truncation mutants. CB, CNOT1 binding domain; P/Q/N, P/Q/N rich aggregation-prone region; YTH, m6A RNA binding domain **B.** HEK293T cells were co-transfected with full length YTHDF2 and truncated YTHDF2 plasmids as indicated. WB analyses showing that PIAS1 is Co-IPed with N-terminal and C-terminal regions of YTHDF2. β-actin blot was included as loading controls. Arrows denoted the position of full-length and truncated YTHDF2. **C.** The schematic representation showing PIAS1 domains, V5-tagged or Halo-V5 tagged PIAS1 truncation mutants. SAP (SAF-A/B, Acinus, and PIAS): DNA and protein binding domain; PINIT: nuclear localization motif; RING: RING finger E3 ligase domain for protein SUMOylation; ST Rich: variable Ser/Thr rich region. **D.** HEK293T cells were co-transfected with HA-YTHDF2 and truncated versions of V5-PIAS1 or Halo-V5-PIAS1. WB analysis showing that YTHDF2 is Co-IPed with the N-terminal and middle part of PIAS1. β-actin blot was included as loading controls. **E.** A proposed model showing the N- and C-terminal regions of YTHDF2 binding to the central part of PIAS1.

To determine the region(s) within PIAS1 that interacts with YTHDF2, we co-transfected several truncated PIAS1 mutants with YTHDF2 into HEK293T cells (**Figure 2C**). We then IPed YTHDF2 with anti-HA antibody conjugated beads. We found that PIAS1 (aa 1-433), PIAS1 (aa 101-433), PIAS1 (aa 1-415) and PIAS1 (aa 1-205), but not PIAS1 (aa 409-651) and PIAS1 (aa 1-100), are Co-IPed by YTHDF2, suggesting that the central region of PIAS1 between SAP and RING domains (aa 101-205) is essential for YTHDF2 binding (**Figure 2D**).

Together, our results indicated that both N-terminal and C-terminal parts of YTHDF2 interact with PIAS1, specifically with the middle region of PIAS1 (**Figure 2E**).

### PIAS1 promotes the anti-EBV activity of YTHDF2

To investigate the potential function of PIAS1 on the anti-viral activity of YTHDF2, we performed co-transfection experiments. We transfected plasmids encoding ZTA (a trigger for EBV lytic reactivation), YTHDF2, and either full-length or truncated forms of PIAS1 into HEK293 (EBVLJ+) cells. We observed that transfection of YTHDF2 alone led to a partial reduction in EBV lytic replication while co-transfection of PIAS1 and YTHDF2 resulted in a significant reduction of EBV DNA replication (**Figure 3**, lane 2 vs 3 and lane 3 vs 4). Interestingly, when truncated forms of PIAS1 were co-transfected, the enhanced anti-EBV activity of YTHDF2 by PIAS1 was compromised, despite some of the fragments still exhibiting binding to YTHDF2 (**Figure 3**, lane 4 vs 5-7). Together, these results suggested that PIAS1 promotes the anti-EBV activity of YTHDF2.

**Figure 3.**
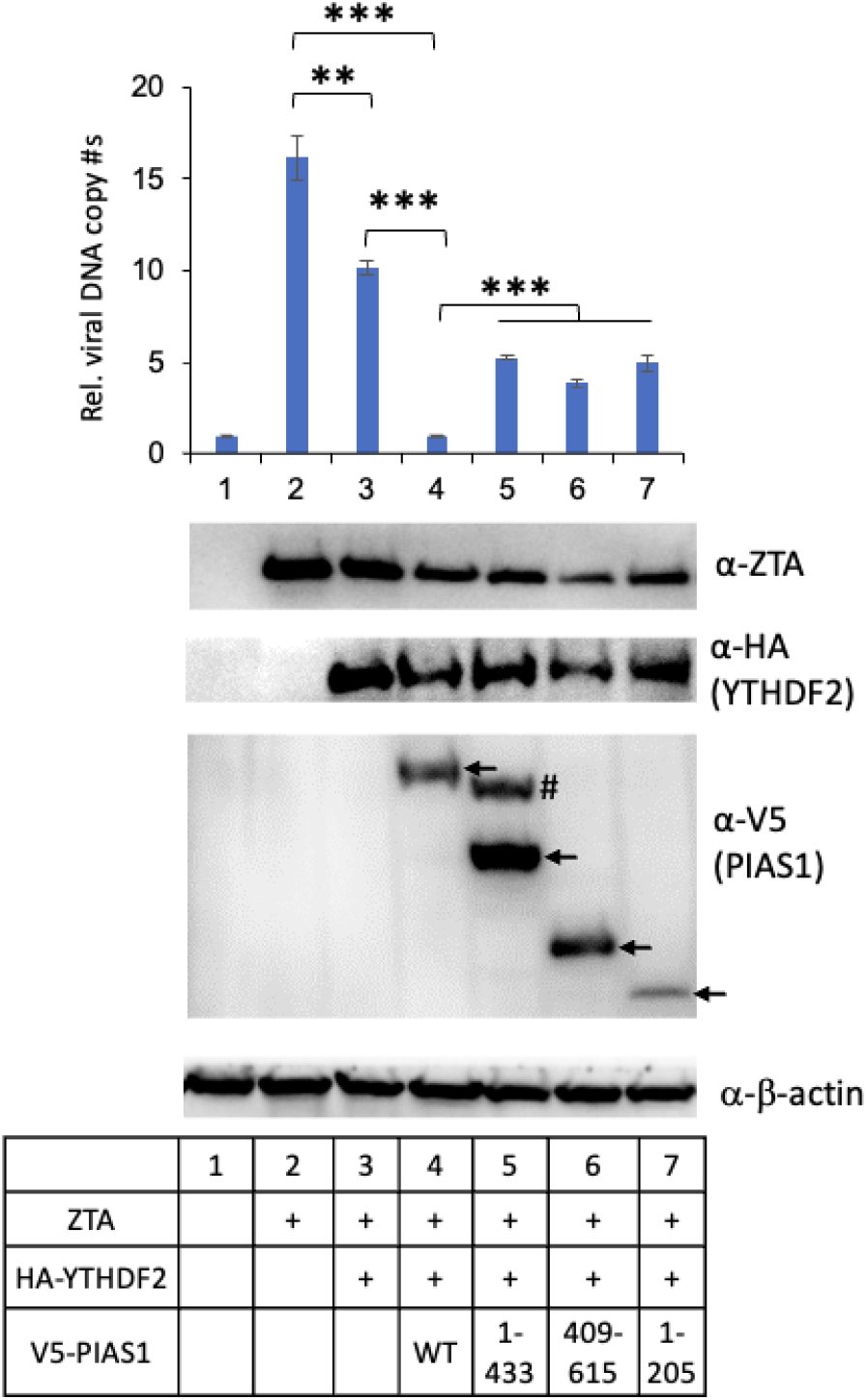
PIAS1 enhances the anti-EBV activity of YTHDF2. HEK293 (EBV+) cells were co-transfected with plasmids encoding ZTA, YTHDF2 and truncated forms of PIAS1 (1-433, 409-615, 1-205) as indicated. The relative EBV copy numbers were measured using the qPCR. The value of lane 1 was set as 1. The protein expression levels were monitored by WB using antibodies as indicated. β-actin blot was included as loading controls. Arrows denote the relative positions of PIAS1 fragments. # denotes a non-specific band. Results from three biological replicates are presented. Error bars indicate standard deviation. **p<0.01, ***p<0.001.

### PIAS1 SUMOylates YTHDF2 at three major sites

The strong interaction between PIAS1 and YTHDF2, along with the enhanced SUMOylation of YTHDF2 in the presence of PIAS1 (**Figures 1-2**), supports the hypothesis that PIAS1 directly SUMOylates YTHDF2. To investigate this, we performed *in vitro* SUMOylation assays using recombinant proteins. We utilized purified V5-YTHDF2 protein and incubated it with E1, E2, SUMO2 protein, and purified PIAS1 protein. Our results clearly demonstrated that PIAS1 significantly enhances the SUMOylation of YTHDF2 (**Figure 4A**, lane 3 vs 1-2).

**Figure 4.**
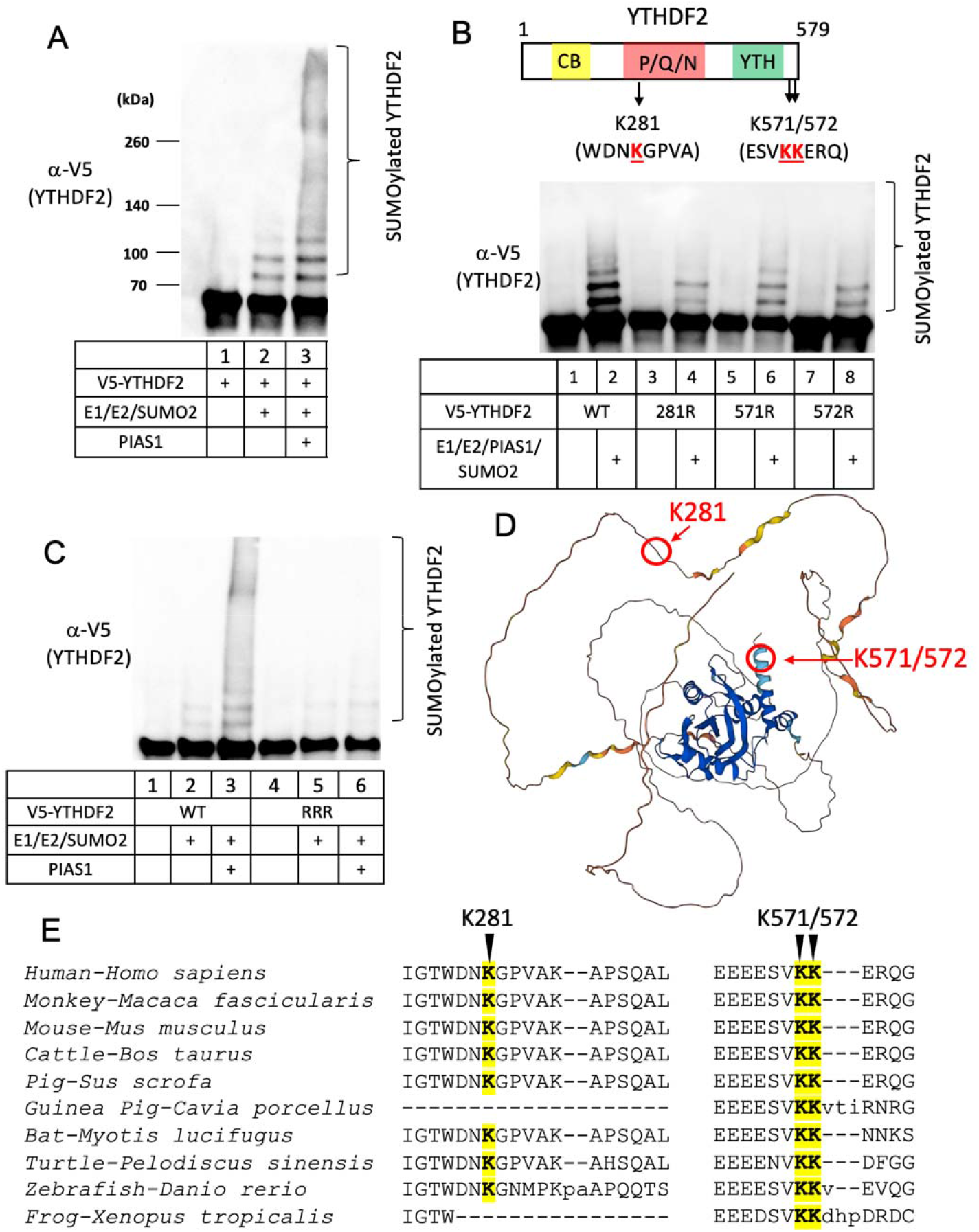
PIAS1 SUMOylates YTHDF2 at K281, K571, and K572. **A.** *In vitro* SUMOylation assay was performed with the combination of purified E1, E2, SUMO2, PIAS1 and substrate V5-YTHDF2 proteins as indicated. The reaction was terminated with SDS sample loading buffer and WB was performed using anti-V5-HRP antibody. Bracket denotes SUMOylated YTHDF2. **B.** The schematic representation of YTHDF2 protein and the corresponding lysine residues (red) within SUMOylation consensus motifs. *In vitro* SUMOylation assay was performed with the combination of E1, E2, SUMO2, PIAS1, and either WT YTHDF2 or individual YTHDF2 mutants (K281R, K571R, and K572R) as indicated. The reaction was terminated with SDS sample loading buffer and WB was performed using anti-V5-HRP antibody. **C.** *In vitro* SUMOylation assay was performed for WT YTHDF2 and RRR mutant (K281R/K571R/K572R) as indicated. **D.** The localization of K281, K571, and K572 in YTHDF2. The YTHDF2 3D structure was predicted by Alphafold. The location of indicated SUMOylation sites were marked by arrows. **E.** Sequence alignment of YTHDF2 protein sequences from 10 species using the Constraint-based Multiple Alignment Tool (COBALT). The SUMOylation sites corresponding to human YTHFD2 were highlighted by yellow color.

Protein SUMOylation typically takes place on lysine residues within the consensus motif ΨKxE/D or the inverted motif E/DxKΨ, where Ψ represents a hydrophobic amino acid and x can be any amino acid residue. However, there are instances where SUMOylation occurs on lysine residues outside of the consensus sequence (17).

To identify the SUMOylation sites on YTHDF2, we searched for the consensus motif and identified two potential SUMOylation sites. The first site is K281, located within the DNKG sequence (DxKΨ), and the second site is K571, located within the VKKE sequence (ΨKxE) (**Figure 4B**, top panel).

To demonstrate whether K281 and K571 can be SUMOylated by PIAS1, we created mutant YTHDF2 constructs in which each lysine residue was individually mutated to arginine. We also created a K572R mutant as it is located within the VKKE sequence. We purified these mutant proteins from HEK293T cells and performed *in vitro* SUMOylation assays.

Our results demonstrated that the SUMOylation level is reduced in all three mutant proteins (K281, K571, and K572) compared to the WT YTHDF2. (**Figure 4B**, bottom panel, lanes 4, 6, 8 vs 2), suggesting that all three sites can be SUMOylated even though K572 is not a consensus SUMOylation site. We then generated a mutant YTHDF2, K281R/K571R/K572R (RRR), in which all three lysine residues were simultaneously mutated to arginines. We found that the SUMOylation of YTHDF2 (RRR) mutant is abolished compared to WT protein (**Figure 4C**, lane 6 vs 3). These findings together demonstrated that K281, K571, and K572 are the major SUMOylation sites on YTHDF2 mediated by PIAS1.

According to the YTHDF2 3-dimensional (3D) structure predicted by Alphafold (18), we observed that K281 is situated within a disordered region, whereas K571 and K572 are located at the C-terminal end of YTHDF2, forming an alpha-helix secondary structure (**Figure 4D**). The presence of a disordered region and the location of lysine residues at the very C-terminal region of YTHDF2 may contribute to structural flexibility favorable for SUMOylation.

To examine the conservation of YTHDF2 SUMOylation sites across different species, we performed an alignment of the amino acid sequence of human YTHDF2 with sequences from nine other species. Remarkably, we observed that the amino acids corresponding to K571/572 of human YTHDF2 are conserved among all the examined species (**Figure 4E**). As for K281 of human YTHDF2, we found that the corresponding sequences are highly conserved in human, monkey, mouse, cattle, pig, bat, turtle, and zebrafish, but are absent in guinea pig and xenopus YTHDF2 (**Figure 4E**). This finding suggested a high likelihood of YTHDF2 SUMOylation by PIAS1 at the same positions in other organisms as observed in humans.

### SUMOylation enhances the anti-EBV activity of YTHDF2

To determine the function of YTHDF2 SUMOylation in EBV replication, we introduced vectors expressing PIAS1, WT YTHDF2 and RRR mutant lacking SUMOylation sites. In the absence of PIAS1 co-transfection, the RRR mutant exhibited higher levels of EBV replication comparted to WT YTHDF2 (**Figure 5A**, lane 5 vs 3), suggesting a reduced anti-viral activity when the SUMOylation of YTHDF2 is blocked. However, when co-transfected with PIAS1, the SUMOylation-deficient mutant displays similar viral replication compared to cells expressing wild-type YTHDF2 (**Figure 5A**, lane 6 vs 4), possibly because PIAS1 overexpression alone also inhibits EBV replication (**Figure 5A**, lane 7).

**Figure 5.**
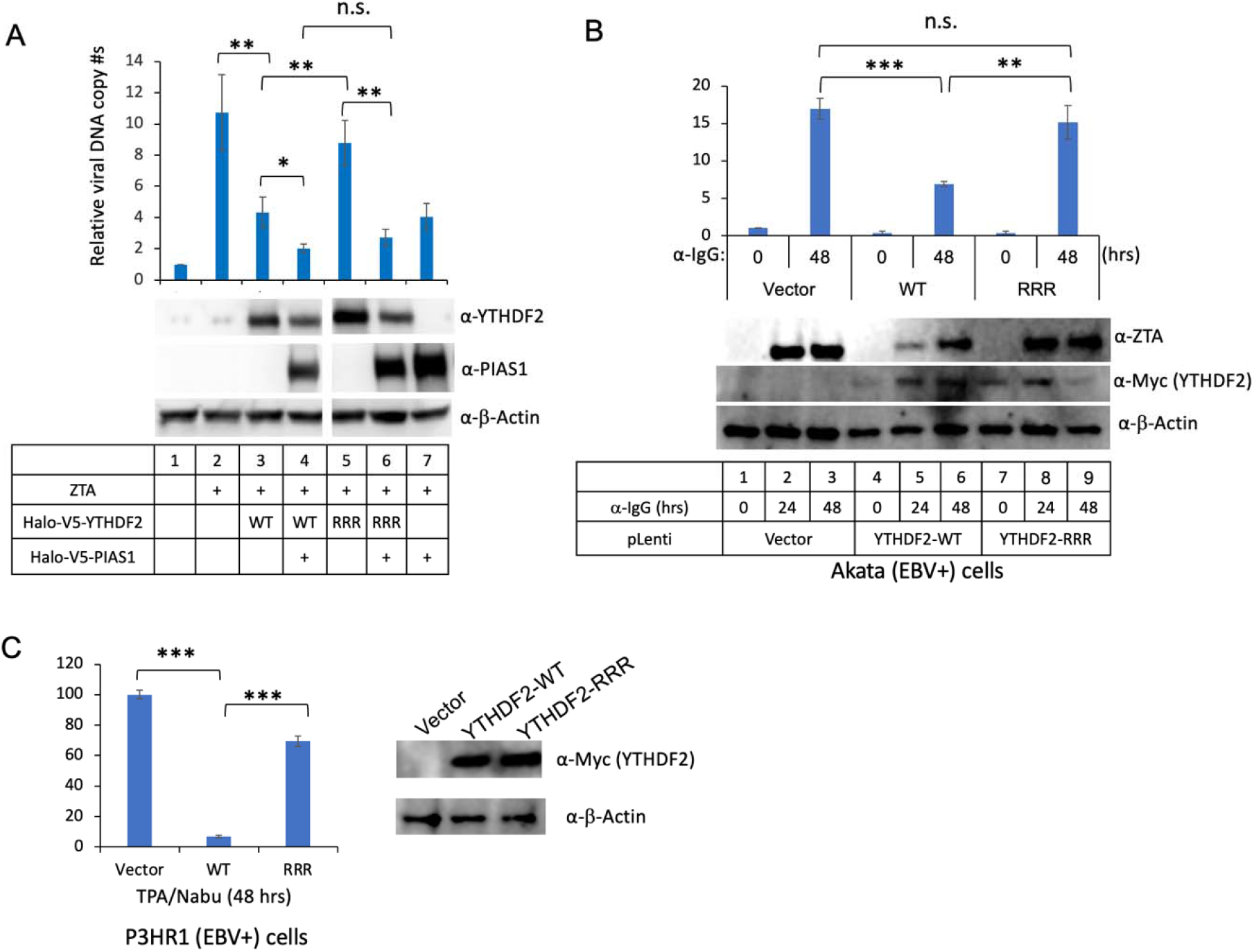
SUMOylation-deficient YTHDF2 impairs its anti-EBV activity. **A.** HEK293 (EBV+) cells were transfected with ZTA plasmid (lytic inducer), PIAS1, and wild-type YTHDF2 (WT) or SUMOylation-deficient YTHDF2 (RRR) as indicated. The relative EBV copy numbers were measured using the qPCR as described in the method. The value of lane 1 was set as 1. The expression levels of YTHDF2 and PIAS1 were monitored by WB. β-actin blot was included as loading controls. **B.** Akata (EBVLJ+) cells were used to create cell lines using pLenti-Vector, pLenti-YTHDF2-WT and p-Lenti-YTHDF2-RRR (K281R/K571R/K572R). EBV lytic cycle was induced by anti-IgG-mediated B cell receptor cross-linking. The relative EBV copy numbers were measured using the qPCR as described in the method. The value of lane 1 was set as 1. The expression levels of ZTA and YTHDF2 were monitored by WB using anti-Myc antibody. β-actin blot was included as loading controls. **C.** P3HR1 (EBVLJ+) cells were used to create cell lines using pLenti-Vector, pLenti-YTHDF2-WT and p-Lenti-YTHDF2-RRR. EBV lytic cycle was induced by TPA and Sodium Butyrate (TPA/NaBu) for 48 hrs. The relative EBV copy numbers were measured using the qPCR as described in the method. The value of lane 1 was set as 100. The expression of YTHDF2 was monitored by WB using anti-Myc antibody. β-actin blot was included as loading controls. Results from three biological replicates are presented. Error bars indicate the standard deviation. *pLJ<LJ0.05; ** pLJ<LJ0.01; ***pLJ<LJ0.001; n.s., not significant.

To demonstrate the physiological relevance of YTHDF2 SUMOylation in EBV replication, we generated lentiviral constructs containing WT and RRR mutant YTHDF2. We transduced Akata (EBV+) Burkitt lymphoma cells with these constructs, establishing stable cell lines expressing either WT or RRR mutant YTHDF2. Upon lytic induction, we found a significant suppression of EBV DNA replication in cells expressing WT YTHDF2 (**Figure 5B**, upper panel). However, we observed that cells expressing the RRR mutant exhibits higher levels of EBV replication compared to those expressing WT YTHDF2 (**Figure 5B**, upper panel). Consistently, we found that viral protein ZTA expression is lower in WT YTHDF2-expressing cells than that in the RRR mutant cells (**Figure 5B**, lower panel, lanes 5 and 6 vs 8 and 9), suggesting that the loss of YTHDF2 SUMOylation abrogates its anti-viral activity.

To further confirm this observation, we also established WT and RRR mutant YTHDF2-expressing cell lines using EBV-positive P3HR1 Burkitt lymphoma cells. Similarly, we found that WT YTHDF2-expressing cells display much lower viral replication than those expressing the RRR mutant upon lytic induction (**Figure 5C**). Together, these findings provided compelling evidence that SUMOylation enhances the anti-EBV activity of YTHDF2.

SUMOylation of YTHDF2 has been implicated in binding to m6A modified RNAs (16). We reasoned that YTHDF2, after SUMOylation, affect its binding to viral lytic genes. To test this possibility, we performed YTHDF2 RNA immunoprecipitation (RIP) assay in Akata (EBV+) cells expressing WT or SUMOylation-deficient YTHDF2. Interestingly we found that SUMOylation-deficient YTHDF2 displays reduced binding to EBV lytic transcripts compared to the WT counterpart. (**Figure 6A**).

**Figure 6.**
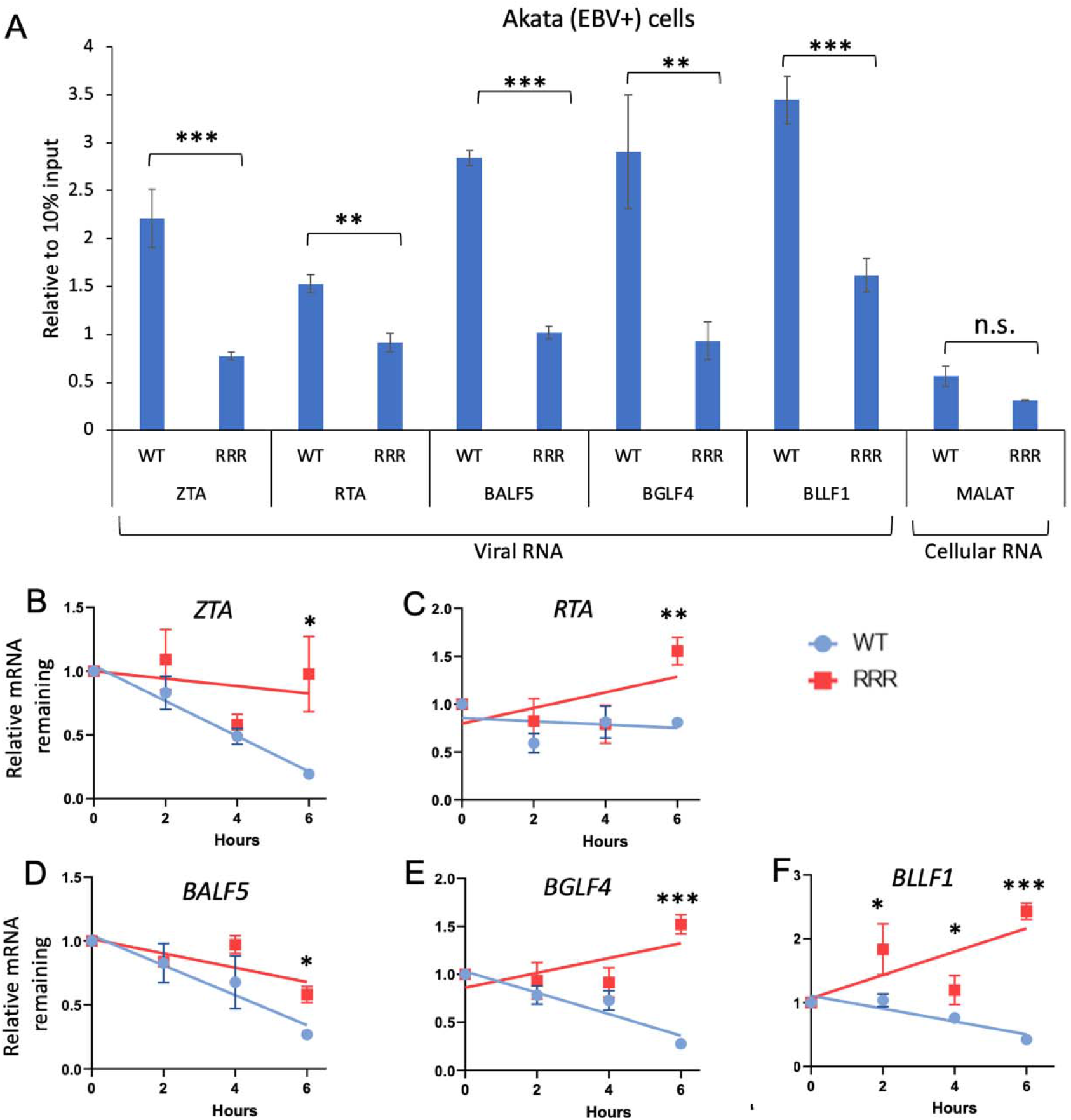
SUMOylation of YTHDF2 enhances its binding to and degradation of EBV lytic transcripts. Akata (EBV+) carrying WT YTHDF2 or SUMOylation-deficient mutant (RRR) were lytically induced by IgG cross-linking for 24 h. **A.** Cell lysate was collected to detect YTHDF2 binding of viral RNAs by RIP-qPCR. Values are fold change over 10% input. Cellular RNA MALAT1 was included a negative control. **B-F.** After lytic induction for 24 h, the cells were then treated with actinomycin D. The Immediate Early (*ZTA* and *RTA*), Early (*BALF5* and *BGLF4*) and Late (*BLLF1*) gene levels were analyzed by qRT-PCR. The relative mRNA level at 0 h after actinomycin D treatment was set as 1. Results from three biological replicates are presented. Error bars indicate SD. *, P< 0.05; **, P< 0.01; ***, P< 0.001, n.s., not significant.

YTHDF2 binding to viral RNA was shown to promote their degradation (5, 7). Therefore, the reduced binding seen in SUMOylation-deficient YTHDF2 may affect RNA stability. To test this idea, we monitored EBV lytic gene stability after actinomycin D treatment of WT or SUMOylation-deficient YTHDF2-expressing Akata (EBV+) cells preinduced with anti-IgG for 24LJh. We found that cells carrying SUMOylation-deficient YTHDF2 displays higher EBV lytic gene stability than cells with WT YTHDF2 (**Figure 6B-F**).

Together, these results suggested that SUMOylated YTHDF2 has higher binding affinity to EBV lytic transcripts, therefore facilitating their decay to restrict EBV lytic replication.

### PIAS1 facilitates the SUMOylation of YTHDF1 and YTHDF3

YTHDF1 and YTHDF3 are paralogs of YTHDF2, and they exhibit redundant functions in regulating mRNA degradation within cells (19). Notably, we and others have demonstrated that both YTHDF1 and YTHDF3 play a role in restricting EBV infection (7, 20). By aligning the amino acid sequences of YTHDF1 and YTHDF3 with that of YTHDF2, we observed that the lysine corresponding to YTHDF2 K281 is highly conserved in both YTHDF1 and YTHDF3. Additionally, while YTHDF2 K571 is not conserved in YTHDF1 and YTHDF3, YTHDF1 doe show conservation of the lysine corresponding to YTHDF2 K572 (**Figure 7A**).

**Figure 7.**
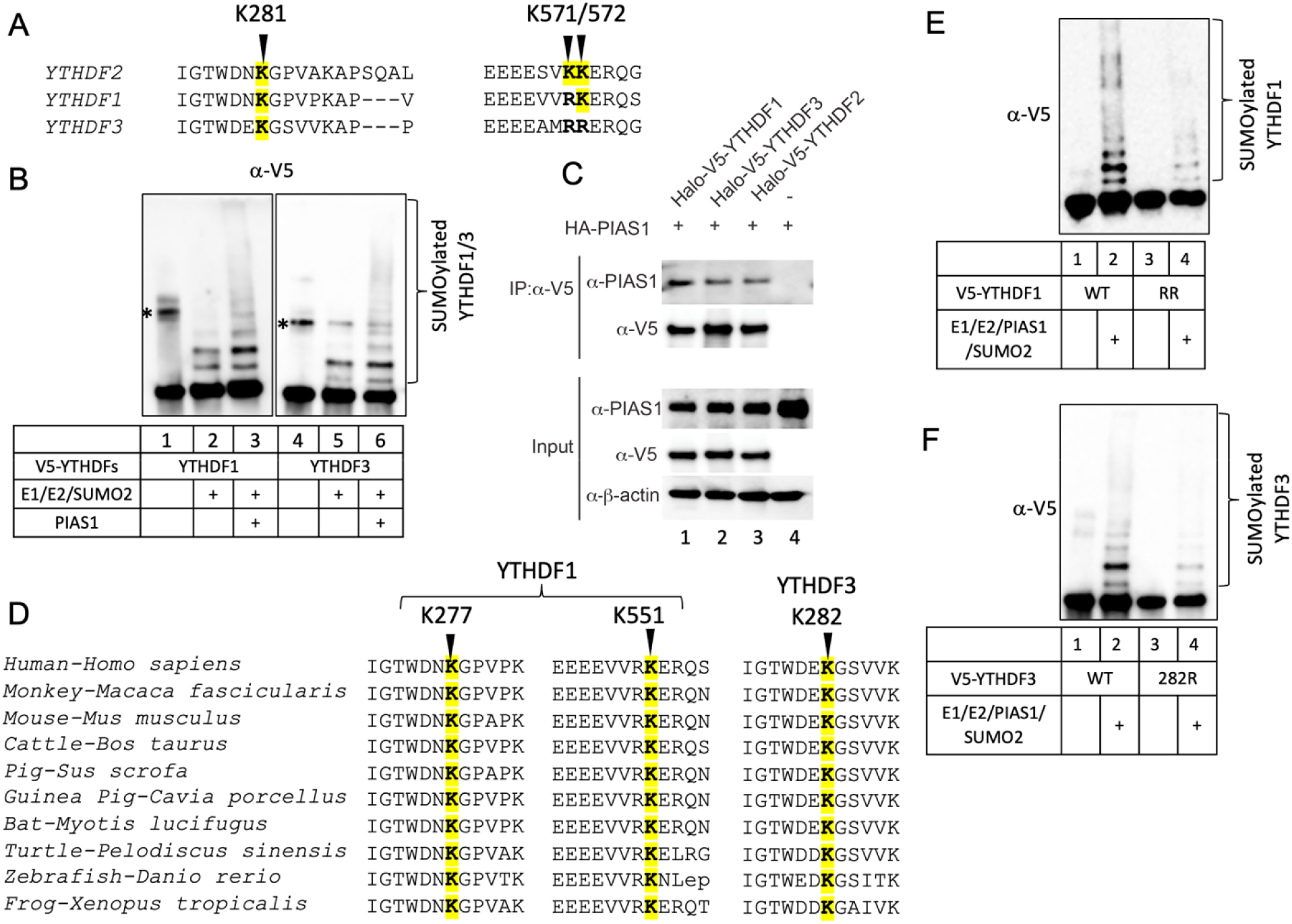
YTHDF1 and YTHDF3 are also SUMOylated by PIAS1. Related to Figure S1. **A.** Sequence alignment of human YTHDF2, YTHDF1 and YTHDF3 using COBALT. The SUMOylation sites corresponding to human YTHDF2 were highlighted by yellow color. **B.** PIAS1 promotes YTHDF1 and YTHDF3 SUMOylation. In vitro SUMOylation of YTHDF1 and YTHDF3 with PIAS1 and WB was performed using anti-V5 antibody. *, non-specific band. **C.** PIAS1 interacts with YTHDF1 and YTHDF3. HEK293T cells were co-transfected with HA-PIAS1 and V5-YTHDF1, V5-YTHDF3, as well as V5-YTHDF2 as indicated. WB analysis showing that PIAS1 is Co-IPed with YTHDF1 and YTHDF3. Whole cell lysates were blotted for PIAS1 and V5-YTHDFs as input. β-actin blot was included as loading controls. **D.** Sequence alignment of YTHDF1 and YTHDF3 protein sequences from 10 species using COBALT. The positions of lysine corresponding to human YTHDF1 and YTHDF3 were labeled as indicated. The conserved lysines were highlighted by yellow color. **E.** *In vitro* SUMOylation assay was performed with the combination of E1, E2, SUMO2, PIAS1, and either WT or K277R/K551R mutant (RR) YTHDF1. The reaction was terminated with SDS sample loading buffer and WB was performed using anti-V5-HRP antibody. **F.** *In vitro* SUMOylation assay was performed with the combination of E1, E2, SUMO2, PIAS1, and either WT or K282R mutant YTHDF3.

Based on this observation, we hypothesized that YTHDF1 and YTHDF3 can undergo SUMOylation and that PIAS1 might promote their SUMOylation. To test this hypothesis, we performed *in vitro* SUMOylation assays using purified YTHDF1 and YTHDF3, along with PIAS1 and the necessary enzymes and proteins (E1, E2, SUMO2). The results from WB analysis demonstrated that YTHDF1 and YTHDF3 are SUMOylated and the presence of PIAS1 enhances the SUMOylation of YTHDF1 and, to a lesser extent, YTHDF3 (**Figure 7B**, lane 3 vs 1-2 and lane 6 vs 4-5).

To investigate the interaction between YTHDF1, YTHDF3 and PIAS1, we transfected HEK293T cells with plasmids expressing HA-PIAS1, V5-YTHDF1, V5-YTHDF3, and V5-YTHDF2 as a control. Following immunoprecipitation of YTHDF proteins using antibodies against V5 tag, we observed a robust interaction between PIAS1 and both YTHDF1 and YTHDF3, similar to the interaction observed with YTHDF2 (**Figure 7C**, lanes 1-2 vs 3).

We analyzed the sequence conservation of YTHDF1 and YTHDF3 with nine other species. The analysis revealed that the lysines corresponding to K277 and K551 in human YTHDF1, as well as K282 in human YTHDF3, are conserved across all the examined species (**Figure 7D**).

To demonstrate whether K277 and K551 in YTHDF1 and K282 in YTHDF3 can be SUMOylated by PIAS1, we created mutant constructs in which lysine residues were mutated to arginines. We purified these mutant proteins from HEK293T cells and performed *in vitro* SUMOylation assays.

We found that the SUMOylation levels of YTHDF1(K277R/K551R) and YTHDF3 (K282R) are greatly reduced compared to their WT counterparts (**Figure 7E and 7F**). These findings together demonstrated that K277 and K551 in YTHDF1 and K282 in YTHDF3 are the major SUMOylation sites mediated by PIAS1. The residual SUMOylation in these mutants also suggested other sites could be SUMOylated.

These results suggested that the targeting of YTHDF family proteins for SUMOylation by PIAS1 is a conserved phenomenon. The presence of conserved residues in these regions implies that SUMOylation of YTHDFs, as well as the targeting by PIAS1, may have important functional implications across different species.

## Discussion

The m6A RNA modification pathway has been implicated in a variety of cellular process by controlling RNA stability, splicing and translation (2, 3, 21). This pathway is regulated by a group of cellular proteins that act as “writers,” “erasers,” and “readers” of the m6A mark. METTL3, METTL14, WTAP, VIRMA and other associated proteins function as writers to methylate the specific adenosines in the RNA molecules. FTO and ALKBH5 function as erasers to remove the methyl group and thereby reversing the modification. The readers, including YTHDF1/2/3, YTHDC1/2 and HNRNPA2B1, are responsible for binding to the m6A-modified RNA to regulate RNA metabolism (2). Recently, the m6A RNA modification pathway has been studied for its role in the life cycle of a variety of viruses, including herpesviruses (21-23). M6A modification and YTHDF2 reader have been shown to regulate KSHV RNA stability and reactivation (4, 5).

We and other have shown that the m6A writers (METTL3, METTL14, WTAP, and VIRMA) and readers (YTHDF1, YTHDF2, and YTHDF3) restrict EBV reactivation (6, 7, 20, 24). On the other hand, several studies suggested that the m6A RNA modification enzymes and readers regulate interferon production to control the infection of KSHV, EBV, human cytomegalovirus (HCMV), vaccinia virus (VACV), and herpes simplex virus type 1 (HSV-1) (25-30).

PTMs, such as ubiquitination, SUMOylation, phosphorylation, methylation, acetylation and proteolytic cleavage, play crucial roles in determining the fate and function of proteins. Several members of the m6A RNA modification pathway undergo PTMs. We recently have shown that caspase-mediated cleavage of the m6A pathway writers and readers promotes viral replication (7). Phosphorylation has been implicated in the regulation of the m6A methyltransferase complex (31). Although WTAP and METTL3 phosphorylation has been shown to enhance the activity of the methyltransferase complex through protein stabilization (32), alphaherpesvirus pseudorabies virus (PRV) and HSV-1 encoded protein kinases US3 inactivate the writer complex through phosphorylation (33). While YTHDF2 has been shown to be degraded via ubiquitin-proteasome pathway when cyclin-dependent kinase 1 (CDK1) is activated (10), phosphorylation of YTHDF2 by EGFR/SRC/ERK signaling prevents its degradation (9).

The SUMOylation of METTL3 has been demonstrated to slightly suppress its methyltransferase activity (Du et al., 2018). ALKBH5 is SUMOylated by PIAS4 to inhibit its m6A demethylase activity by blocking substrate accessibility (34). Additionally, YTHDF2 has also been identified as a target for SUMOylation, which enhances its binding affinity to m6A-modified RNAs and promotes their degradation (Hou et al., 2021).

In this study, we for the first time demonstrated that YTHDF2 is SUMOylated by the E3 SUMO ligase PIAS1 (**Figures 1** and **4**). We observed that both the N-terminal and C-terminal regions of YTHDF2 are involved in the interaction with PIAS1. Similarly, the central part of PIAS1 was found to interact with YTHDF2 (**Figure 2**). The strong interaction between YTHDF2 and PIAS1 also suggested that PIAS1 may regulate the anti-viral activity of YTHDF2. Indeed, we found that the anti-EBV activity of YTHDF2 was significantly enhanced in the presence of full-length PIAS1, but less so in the presence of truncated PIAS1 mutants (**Figure 3**). These results suggested that the regulation of YTHDF2 SUMOylation mediated by PIAS1 contributes to its anti-viral activity, like our previous observations with SAMHD1 (14).

During our investigation into the SUMOylation of YTHDF2, we remained open-minded regarding the potential presence of additional SUMOylation sites, despite the identification of K571 as one modification site in a previous study (16). Interestingly, we uncovered two novel SUMOylation sites in YTHDF2, namely K281 and K572 (**Figure 4**).

The discovery of K281 as a SUMOylation site in YTHDF2 highlights a modification occurring within the P/Q/N disordered region, which is highly conserved among YTHDF2 homologs from various species (**Figure 4**). The identification of K572 reveals the presence of a non-consensus SUMOylation site that may have a redundant role in YTHDF2 SUMOylation, which further supports the notion that non-consensus SUMOylation is a common event in cells (17, 35).

To investigate the role of YTHDF2 SUMOylation in EBV replication, we generated a SUMOylation-deficient YTHDF2 mutant. Our findings revealed SUMOylation-deficient YTHDF2 displayed reduced anti-viral activity due to its reduced binding to and degradation of viral RNAs (**Figure 6**) (16). Moreover, the establishment of stable cell lines expressing WT or mutant YTHDF2 further stressed the importance of SUMOylation in its anti-EBV activity in multiple EBV-infected tumor cells (**Figure 5**). Notably, previous studies have demonstrated that YTHDF2 undergoes ubiquitination (10, 30). Therefore, SUMOylation could potentially hinder ubiquitination and subsequent degradation of YTHDF2, which requires further investigation.

The sequence conservation between YTHDF2 and its paralogs led us to investigate the potential SUMOylation of YTHDF1 and YTHDF3 by PIAS1 (**Figure 7**). We found that the amino acids corresponding to K281 in YTHDF2, namely K277 in YTHDF1 and K282 in YTHDF3, are conserved across all ten species examined. Consistently, the mutation of these sites reduced their SUMOylation level by PIAS1. The residual SUMOylation suggested additional sites could b SUMOylated. PIAS1 interacts with all members of the YTHDF family. Interestingly, structur prediction by a recently developed AlphaFold-Multimer algorithm (36) suggested that the PIAS1 aa 161-239 region interacts with the YTH domain of YTHDF1, YTHDF2 and YTHDF3 (**Figure S1**). This is consistent with our Co-IP results showing that the central part of PIAS1 (aa 101-433) interacts with YTHDF2 (**Figure 2**).

In conclusion, our findings demonstrated that YTHDF2 undergoes SUMOylation at three key sites, and PIAS1 plays a crucial role in promoting the anti-EBV activity of YTHDF2 by enhancing its SUMOylation (**Figure 8**). Our observations provided insights into the conservation and potential functional significance of PIAS1-mediated SUMOylation of YTHDF2, as well as YTHDF1, and YTHDF3, suggesting that SUMOylation may play a role in regulating the activities of these proteins in RNA metabolism and consequently other cellular processes. METTL3, as demonstrated by Du et al. in 2018, has already been identified as a target for regulation through SUMOylation (37). It will be interesting to explore whether METTL3 and other members of the m6A RNA modification pathway are regulated by PIAS1-mediated SUMOylation and the associated impacts on RNA modification, viral infection and host defense.

**Figure 8.**
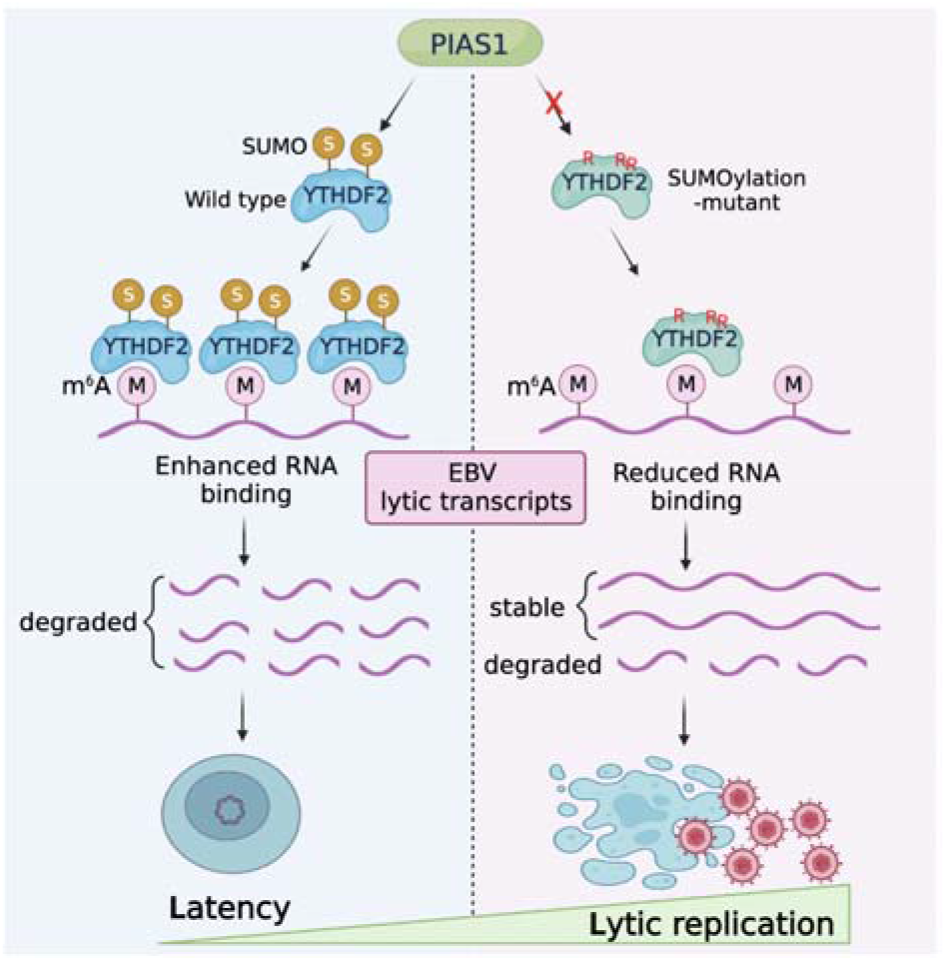
Model summarizing PIAS1-mediated SUMOylation of YTHDF2 in control of EBV latency and reactivation.

## Material and Methods

**Table 1.**
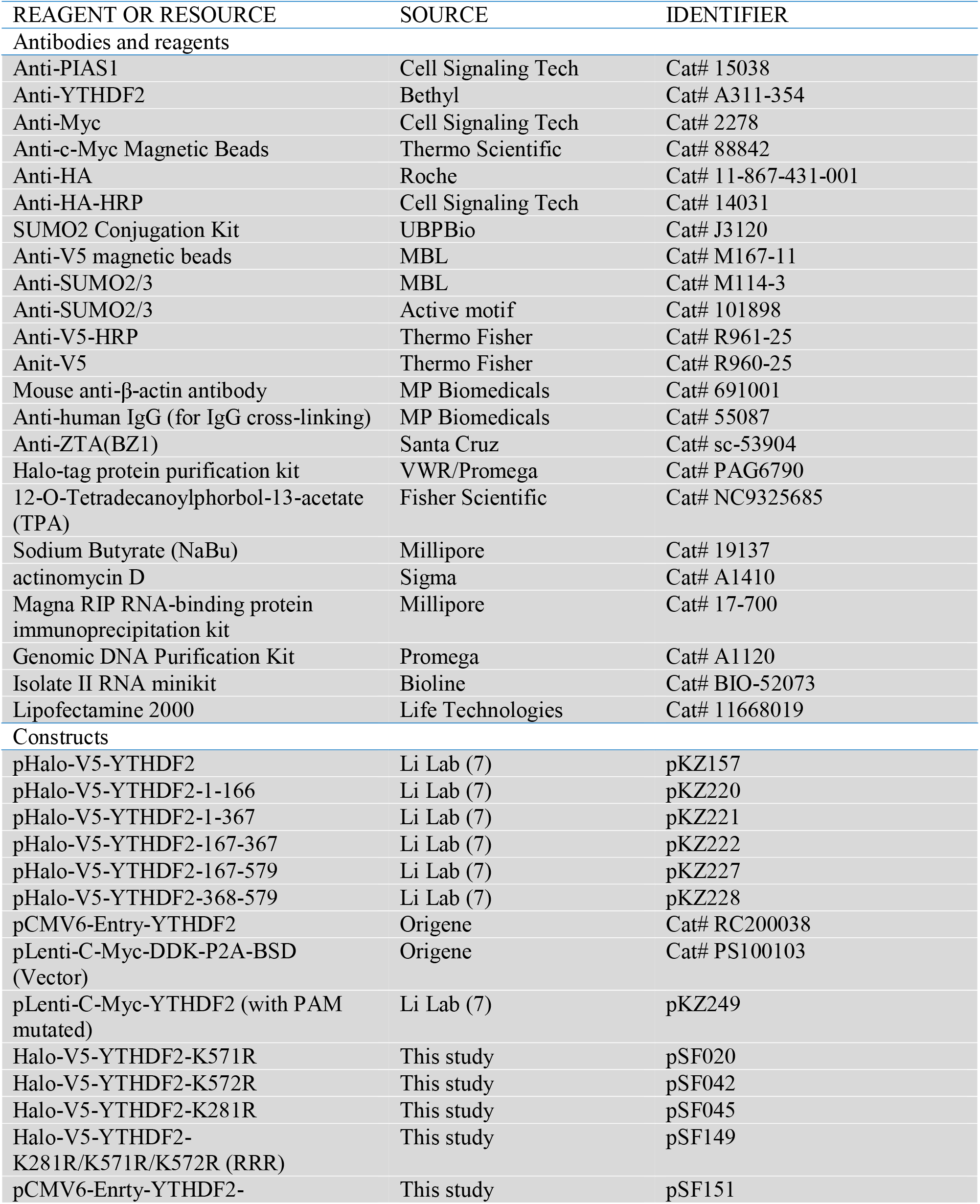
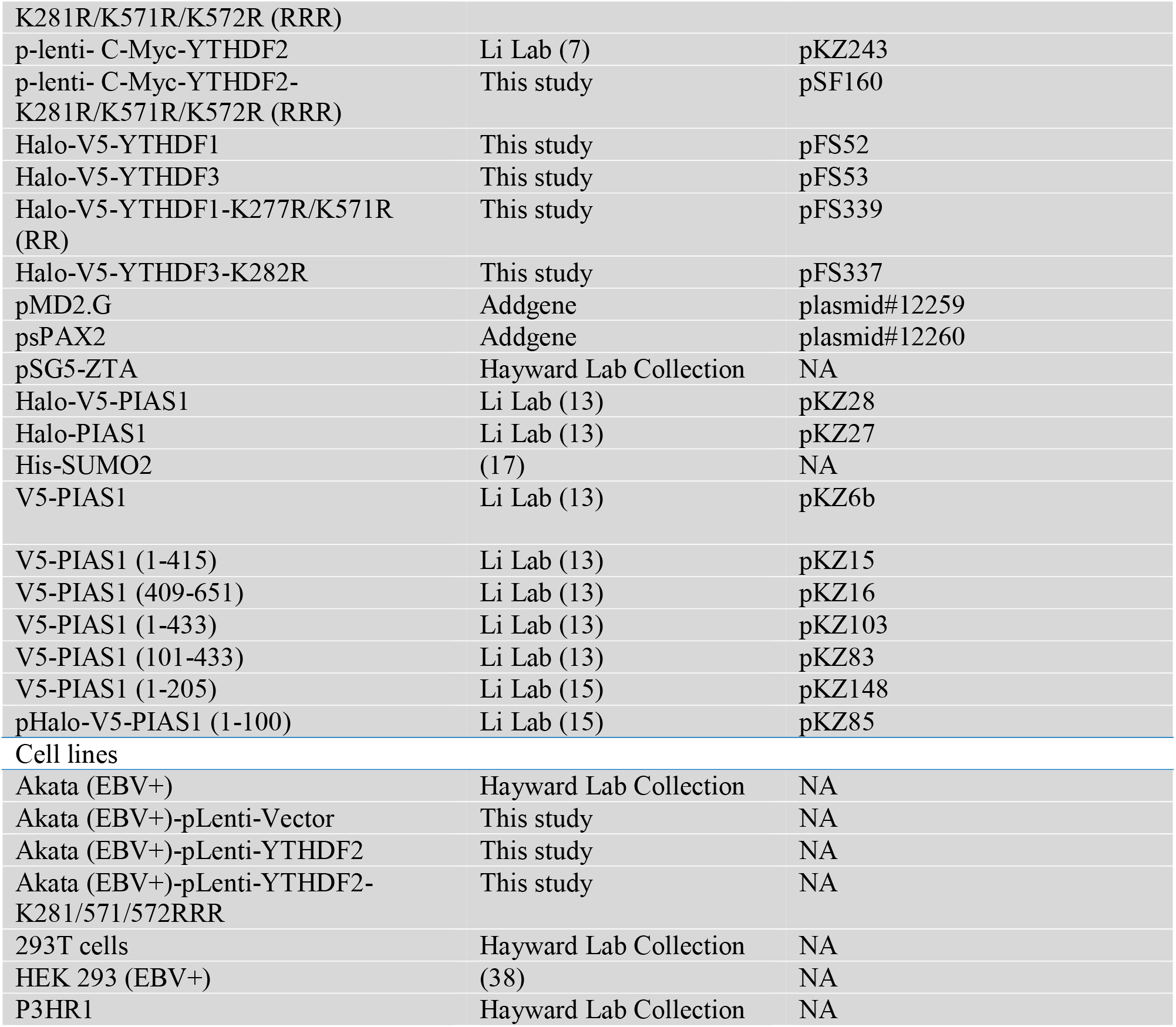
Key reagents and resources.

### Cell Lines and Cultures

Akata (EBV+) and P3HR-1 cells were cultured in Roswell Park Memorial Institute medium (RPMI 1640) supplemented with 10% FBS (Cat# 26140079, Thermo Fisher Scientific) in 5% CO_2_ at 37 °C (13, 39-42). HEK293 (EBV+) cells with B95.8 EBV genome was provided by Henri-Jacques Delecluse and Ayman EI-Guindy (38, 43). HEK293 (EBV+) and 293T cells were cultured in Dulbecco’s modified Eagle medium (DMEM) supplemented with 10% FBS in 5% CO_2_ at 37 °C.

### Plasmids Construction

Halo-PIAS1, Halo-V5-PIAS1 (full length and aa 1-100), and V5-PIAS1 (full length, aa 1-205, aa 1-415, aa 1-433, aa 409-651, and aa 101-433) plasmids were previously described (13, 15). Halo-V5-YTHDF2, (full length, aa 1-166, aa 1-367, aa 167-367, aa 167-579, and aa 368-579) plasmids were constructed by our lab previously (7). Halo-V5-YTHDF2 was used as template to create K281R, K571R, and K572R mutants using the QuikChange II site-Directed Mutagenesis Kit (Cat # 200521; Stratagene) according to the manufacturer’s instructions. Halo-V5-YTHDF1 and Halo-V5-YTHDF3 were constructed as using methods as previously described (7). These plasmids were then used to create Halo-V5-YTHDF1(K277R/K552R) and Halo-V5-YTHDF3 (K282R).

Halo-V5-YTHDF2-K281R/K571R/K572R was created using QuikChange II site-Directed Mutagenesis Kit step by step. The pLenti-C-Myc-DDK-P2A-BSD and pCMV6-Entry-YTHDF2 were purchased from Origene. The specific variants were generated by site-directed mutagenesis in pCMV6-Entry-YTHDF2 using the QuikChange II site-directed mutagenesis kit. Subsequently, the WT and mutated YTHDF2 in pCMV6-Entry vector were digested using AsiSI and MluI and subcloned into the pLenti-C-Myc-DDK-P2A-BSD vector. All primer sequences are listed in **Table S1**.

### Lentiviral Transduction

Lentiviruses were prepared using 293T cells transfected with lentiviral vector containing the target gene and the helper vectors (pMD2.G and psPAX2) as previously descripted (13, 44). Akata (EBV+) and P3HR-1 cells were transduced with lentiviruses carrying vector control, WT and mutant YTHDF2 to establish cell lines in RPMI medium containing 2 μg/ml puromycin.

### Cell Lysis, Immunoprecipitation and Immunoblotting

Cells lysis, immunoprecipitation and immunoblotting (Western Blotting, WB) were performed as previously descripted (14).

### Protein Expression and Purification

Halo-tagged PIAS1, YTHDF2, YTHDF1, and YTHDF3 proteins were expressed and purified as previously described (13, 41, 44).

### *In Vitro* SUMOylation Assay

In vitro SUMOylation assay was performed using the SUMO2 Conjugation Kit. The assay was conducted in a buffer comprising 40 mM Tris pH 7.1, 40 mM NaCl, 1 mM β-mercaptoethanol (ME), and 5 mM MgCl_2_. Each protein SUMOylation reaction included 100 nM SAE1/SAE2 (E1), 2 µM 6xHis-Ube2I/UBC9 (E2), 50 µM SUMO2, and 4 mM ATP. Additionally, purified PIAS1 was introduced as the E3 ligase. The reaction mixtures were then incubated at 37LJ°C for 3LJh, and WB analysis was carried out to examine the SUMOylation of YTHDF2, YTHDF1, and YTHDF3 proteins.

### Lytic Induction

Akata (EBV+) cells were treated with IgG (1:200, Cat# 55087, MP Biomedicals) to induce lytic replication for up to 48 hrs. To induce the EBV lytic cycle in P3HR-1 cells, the cells were triggered with tetradecanoyl phorbol acetate (TPA; 20 ng/ml) and sodium butyrate (3 mM) for up to 48LJhrs (7). For lytic induction of EBV in HEK293 (EBV+) cells, the cells were transfected with EBV ZTA plus other plasmids as appropriate using Lipofectamine 2000 reagent for 48 hrs.

### EBV Copy Number Detection

To extract cell associated viral DNA, total genomic DNA was extracted using the Genomic DNA Purification Kit (Cat# A1120, Promega). The relative viral genome copy numbers were determined by quantitative polymerase chain reaction (qPCR) using primers specific to *BALF5* gene normalized by *β-actin* as we described previously (44). Extracellular viral DNA was extracted and measured as previously described (42).

### RNA-binding protein immunoprecipitation

RNA-binding protein immunoprecipitation (RIP) was conducted using the Magna RIP kit (catalog no. 17-700; Millipore) following the manufacturer’s instructions. In brief, Akata (EBV+) cells expressing YTHDF2-Myc or YTHDF2 (K281R/K571R/K572R, RRR)-Myc were treated with IgG cross-linking for 24 hours and then lysed using the RIP lysis buffer provided in the kit. A portion of the lysate (10%) was saved as the input sample. The anti-c-Myc magnetic bead (catalog no. 88842, Thermo Scientific) were washed with RIP buffer and incubated with RNA overnight at 4°C. On the following day, the beads were collected and washed six times with the RIP wash buffer. The enriched RNA-protein complex was digested with proteinase K, and the released RNA was purified using phenol-chloroform extraction. The purified RNA was then subjected to reverse transcription for subsequent qPCR analysis, using the primers for *ZTA, RTA BALF5, BGLF4, BLLF1,* and *MALAT1* as we previously described (7, 13)

### mRNA stability assay

Akata (EBV+) cells expressing YTHDF2-Myc or YTHDF2 (K281R/K571R/K572R, RRR)-Myc in 6-well plates were treated with IgG cross-linking for 24LJhrs. The cells where then treated with actinomycin D (5 μg/ml) (catalog no. A1410; Sigma-Aldrich) to inhibit transcription. The cells were collected at 0, 2, 4, and 6LJhrs after treatment. The total RNA was extracted with an Isolate II RNA minikit (Bioline) and analyzed by qRT-PCR with specific primers for *ZTA, RTA, BALF5, BGLF4,* and *BLLF1*. 18s rRNA was used as control (45). All primer sequences are listed in **Table S1**.

### Structure prediction by AlphaFold-Multimer

AlphaFold-Multimer algorithm (36) was employed to predict protein-protein interactions involving the following sequences: PIAS1 (Uniprot: O75925), YTHDF1 (Uniprot: Q9BYJ9), YTHDF2 (Uniprot: Q9Y5A9), and YTHDF3 (Uniprot: Q7Z739). The detailed procedure was described in the link: https://cosmic-cryoem.org/tools/alphafoldmultimer/ (46, 47). Briefly, the amino acid sequences of PIAS1 and YTHDF1, YTHDF2 or YTHDF3 were combined into fasta files, which were then uploaded to the COSMIC2 webserver. This fasta file was used as the input to run the AlphaFold-Multimer tool. Each prediction of protein-protein interaction generated 5 pdb files. Molecular graphics and analyses of protein interactions were performed with UCSF ChimeraX (48), developed by the Resource for Biocomputing, Visualization, and Informatics at the University of California, San Francisco, with support from National Institutes of Health R01-GM129325 and the Office of Cyber Infrastructure and Computational Biology, National Institute of Allergy and Infectious Diseases. Model 1 of each prediction was used to display PIAS1 interaction with YTHDF1, YTHDF2 and YTHDF3.

### Quantification and Statistical Analysis

Statistical analyses were performed using a two-tailed Student’s t test with Microsoft Excel software. A p value less than 0.05 was considered statistically significant. The values are presented as means and standard deviations (SD) for biological replicate experiments as specified in the figure legends. For RNA decay assay, the figures were created using GraphPad Prism 9 software. The Figure 8 was created using BioRender.

## Acknowledgements

We extend our gratitude to S. Diane Hayward from Johns Hopkins University for providing the reagents and cell lines. We would also like to thank Didier Trono from EPFL for providing the pMD2.G and psPAX2 plasmids (Addgene plasmid #s 12259 and 12260) and Ronald Hay from University of Dundee for providing His-SUMO2 plasmid. Additionally, we express our appreciation to Henri-Jacques Delecluse from the German Cancer Research Centre and Ayman El-Guindy from Yale University for providing the HEK293 cells carrying B95.8 EBV genomes. This work was supported, in part, from grants from the National Institute of Allergy and Infectious Diseases (AI104828 and AI141410; https://grants.nih.gov/grants/oer.htm) to RL. RL was supported by a Research Scholar Grant (134703-RSG-20–054-01-MPC) from the American Cancer Society. RL also received support from the University of Pittsburgh Medical Center (UPMC) Hillman Cancer Center, Virginia Commonwealth University (VCU) Philips Institute for Oral Health Research, and the VCU Presidential Quest for Distinction Award. The funders had no role in study design, data collection and analysis, decision to publish, or preparation of the manuscript.

## Author Contributions

Conceptualization, RL, FS and FGS

Data curation, FS, FGS, KZ and RL

Formal analysis, FS, FGS, KZ and RL

Funding acquisition, RL

Investigation, FS, FGS, KZ and RL

Methodology, FS, FGS, KZ and RL

Project administration, RL

Resources, RL

Supervision, RL Validation, FS, FGS, KZ and RL

Visualization, FS, FGS, KZ and RL

Writing - original draft, FGS Writing - review & editing, RL, FS, FGS and KZ

---

**Figure S1.**
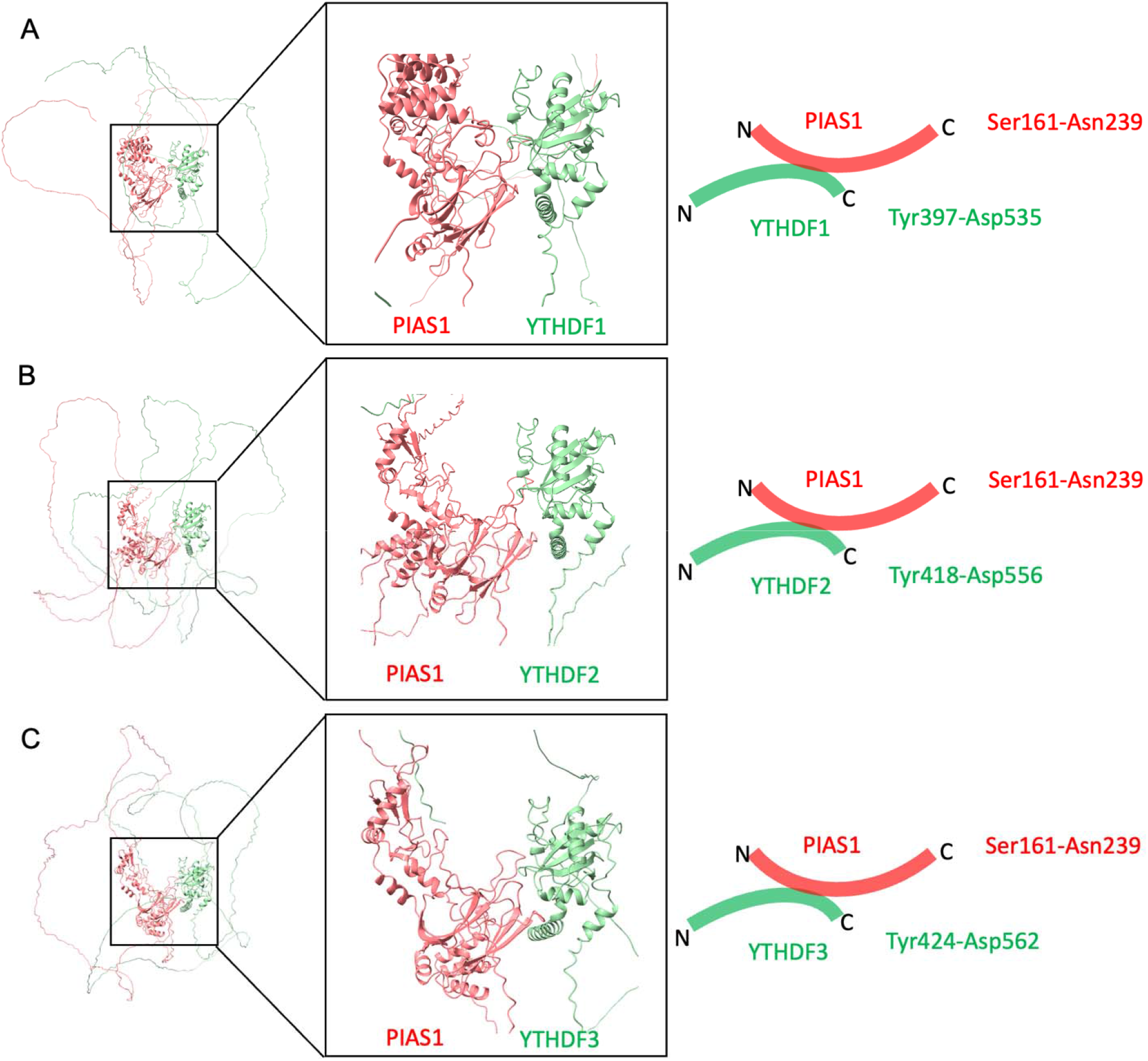
Related to Figures 2 and 7. The AlphaFold-Multimer algorithm was utilized to predict the interaction between PIAS1 and YTHDF1 (A), YTHDF2 (B), and YTHDF3 (C). Among the five models generated, one was selected to visualize the co-structure using the ChimeraX tool. The YTH domains were adjusted to be in similar positions to illustrate their bindings to PIAS1.The amino acids sequences involve in the protein-protein interactions were illustrated.

## References

1. Duggal NK, Emerman M. 2012. Evolutionary conflicts between viruses and restriction factors shape immunity. Nat Rev Immunol 12:687–95.

2. Meyer KD, Jaffrey SR. 2017. Rethinking m(6)A Readers, Writers, and Erasers. Annu Rev Cell Dev Biol 33:319–342.

3. Roundtree IA, Evans ME, Pan T, He C. 2017. Dynamic RNA Modifications in Gene Expression Regulation. Cell 169:1187–1200.

4. Hesser CR, Karijolich J, Dominissini D, He C, Glaunsinger BA. 2018. N6-methyladenosine modification and the YTHDF2 reader protein play cell type specific roles in lytic viral gene expression during Kaposi’s sarcoma-associated herpesvirus infection. PLoS Pathog 14:e1006995.

5. Tan B, Liu H, Zhang S, da Silva SR, Zhang L, Meng J, Cui X, Yuan H, Sorel O, Zhang SW, Huang Y, Gao SJ. 2018. Viral and cellular N(6)-methyladenosine and N(6),2’-O-dimethyladenosine epitranscriptomes in the KSHV life cycle. Nat Microbiol 3:108–120.

6. Lang F, Singh RK, Pei Y, Zhang S, Sun K, Robertson ES. 2019. EBV epitranscriptome reprogramming by METTL14 is critical for viral-associated tumorigenesis. PLoS Pathog 15:e1007796.

7. Zhang K, Zhang Y, Maharjan Y, Sugiokto FG, Wan J, Li R. 2021. Caspases Switch off the m(6)A RNA Modification Pathway to Foster the Replication of a Ubiquitous Human Tumor Virus. mBio 12:e0170621.

8. Imam H, Kim GW, Mir SA, Khan M, Siddiqui A. 2020. Interferon-stimulated gene 20 (ISG20) selectively degrades N6-methyladenosine modified Hepatitis B Virus transcripts. PLoS Pathog 16:e1008338.

9. Fang R, Chen X, Zhang S, Shi H, Ye Y, Shi H, Zou Z, Li P, Guo Q, Ma L, He C, Huang S. 2021. EGFR/SRC/ERK-stabilized YTHDF2 promotes cholesterol dysregulation and invasive growth of glioblastoma. Nat Commun 12:177.

10. Fei Q, Zou Z, Roundtree IA, Sun HL, He C. 2020. YTHDF2 promotes mitotic entry and is regulated by cell cycle mediators. PLoS Biol 18:e3000664.

11. Gareau JR, Lima CD. 2010. The SUMO pathway: emerging mechanisms that shape specificity, conjugation and recognition. Nat Rev Mol Cell Biol 11:861–71.

12. Shuai K, Liu B. 2005. Regulation of gene-activation pathways by PIAS proteins in the immune system. Nat Rev Immunol 5:593–605.

13. Zhang K, Lv DW, Li R. 2017. B Cell Receptor Activation and Chemical Induction Trigger Caspase-Mediated Cleavage of PIAS1 to Facilitate Epstein-Barr Virus Reactivation. Cell Rep 21:3445–3457.

14. Saiada F, Zhang K, Li R. 2021. PIAS1 potentiates the anti-EBV activity of SAMHD1 through SUMOylation. Cell Biosci 11:127.

15. Zhang K, Lv DW, Li R. 2020. Protein inhibitor of activated STAT1 (PIAS1) inhibits IRF8 activation of Epstein-Barr virus lytic gene expression. Virology 540:75–87.

16. Hou G, Zhao X, Li L, Yang Q, Liu X, Huang C, Lu R, Chen R, Wang Y, Jiang B, Yu J. 2021. SUMOylation of YTHDF2 promotes mRNA degradation and cancer progression by increasing its binding affinity with m6A-modified mRNAs. Nucleic Acids Res 49:2859–2877.

17. Tammsalu T, Matic I, Jaffray EG, Ibrahim AFM, Tatham MH, Hay RT. 2014. Proteome-wide identification of SUMO2 modification sites. Sci Signal 7:rs2.

18. Jumper J, Evans R, Pritzel A, Green T, Figurnov M, Ronneberger O, Tunyasuvunakool K, Bates R, Zidek A, Potapenko A, Bridgland A, Meyer C, Kohl SAA, Ballard AJ, Cowie A, Romera-Paredes B, Nikolov S, Jain R, Adler J, Back T, Petersen S, Reiman D, Clancy E, Zielinski M, Steinegger M, Pacholska M, Berghammer T, Bodenstein S, Silver D, Vinyals O, Senior AW, Kavukcuoglu K, Kohli P, Hassabis D. 2021. Highly accurate protein structure prediction with AlphaFold. Nature 596:583–589.

19. Zaccara S, Jaffrey SR. 2020. A Unified Model for the Function of YTHDF Proteins in Regulating m(6)A-Modified mRNA. Cell 181:1582–1595 e18.

20. Xia TL, Li X, Wang X, Zhu YJ, Zhang H, Cheng W, Chen ML, Ye Y, Li Y, Zhang A, Dai DL, Zhu QY, Yuan L, Zheng J, Huang H, Chen SQ, Xiao ZW, Wang HB, Roy G, Zhong Q, Lin D, Zeng YX, Wang J, Zhao B, Gewurz BE, Chen J, Zuo Z, Zeng MS. 2021. N(6)-methyladenosine-binding protein YTHDF1 suppresses EBV replication and promotes EBV RNA decay. EMBO Rep 22:e50128.

21. Tan B, Gao SJ. 2018. RNA epitranscriptomics: Regulation of infection of RNA and DNA viruses by N(6)-methyladenosine (m(6) A). Rev Med Virol 28:e1983.

22. McFadden MJ, Horner SM. 2021. N(6)-Methyladenosine Regulates Host Responses to Viral Infection. Trends Biochem Sci 46:366–377.

23. McFadden MJ, McIntyre ABR, Mourelatos H, Abell NS, Gokhale NS, Ipas H, Xhemalce B, Mason CE, Horner SM. 2021. Post-transcriptional regulation of antiviral gene expression by N6-methyladenosine. Cell Rep 34:108798.

24. Dai DL, Li X, Wang L, Xie C, Jin Y, Zeng MS, Zuo Z, Xia TL. 2021. Identification of an N6-methyladenosine-mediated positive feedback loop that promotes Epstein-Barr virus infection. J Biol Chem 296:100547.

25. Winkler R, Gillis E, Lasman L, Safra M, Geula S, Soyris C, Nachshon A, Tai-Schmiedel J, Friedman N, Le-Trilling VTK, Trilling M, Mandelboim M, Hanna JH, Schwartz S, Stern-Ginossar N. 2019. m(6)A modification controls the innate immune response to infection by targeting type I interferons. Nat Immunol 20:173–182.

26. Rubio RM, Depledge DP, Bianco C, Thompson L, Mohr I. 2018. RNA m(6) A modification enzymes shape innate responses to DNA by regulating interferon beta. Genes Dev 32:1472–1484.

27. Ma S, Yan J, Barr T, Zhang J, Chen Z, Wang LS, Sun JC, Chen J, Caligiuri MA, Yu J. 2021. The RNA m6A reader YTHDF2 controls NK cell antitumor and antiviral immunity. J Exp Med 218.

28. Hesser CR, Walsh D. 2023. YTHDF2 Is Downregulated in Response to Host Shutoff Induced by DNA Virus Infection and Regulates Interferon-Stimulated Gene Expression. J Virol 97:e0175822.

29. Bose D, Lin X, Gao L, Wei Z, Pei Y, Robertson ES. 2023. Attenuation of IFN signaling due to m(6)A modification of the host epitranscriptome promotes EBV lytic reactivation. J Biomed Sci 30:18.

30. Feng J, Meng W, Chen L, Zhang X, Markazi A, Yuan W, Huang Y, Gao SJ. 2023. N(6)-Methyladenosine and Reader Protein YTHDF2 Enhance the Innate Immune Response by Mediating DUSP1 mRNA Degradation and Activating Mitogen-Activated Protein Kinases during Bacterial and Viral Infections. mBio 14:e0334922.

31. Scholler E, Weichmann F, Treiber T, Ringle S, Treiber N, Flatley A, Feederle R, Bruckmann A, Meister G. 2018. Interactions, localization, and phosphorylation of the m(6)A generating METTL3-METTL14-WTAP complex. RNA 24:499–512.

32. Sun HL, Zhu AC, Gao Y, Terajima H, Fei Q, Liu S, Zhang L, Zhang Z, Harada BT, He YY, Bissonnette MB, Hung MC, He C. 2020. Stabilization of ERK-Phosphorylated METTL3 by USP5 Increases m(6)A Methylation. Mol Cell 80:633–647 e7.

33. Jansens RJJ, Verhamme R, Mirza AH, Olarerin-George A, Van Waesberghe C, Jaffrey SR, Favoreel HW. 2022. Alphaherpesvirus US3 protein-mediated inhibition of the m6A mRNA methyltransferase complex. Cell Rep 40:111107.

34. Yu F, Wei J, Cui X, Yu C, Ni W, Bungert J, Wu L, He C, Qian Z. 2021. Post-translational modification of RNA m6A demethylase ALKBH5 regulates ROS-induced DNA damage response. Nucleic Acids Res 49:5779–5797.

35. Blomster HA, Imanishi SY, Siimes J, Kastu J, Morrice NA, Eriksson JE, Sistonen L. 2010. In vivo identification of sumoylation sites by a signature tag and cysteine-targeted affinity purification. J Biol Chem 285:19324–9.

36. Evans R, O’Neill M, Pritzel A, Antropova N, Senior A, Green T, Žídek A, Bates R, Blackwell S, Yim J, Ronneberger O, Bodenstein S, Zielinski M, Bridgland A, Potapenko A, Cowie A, Tunyasuvunakool K, Jain R, Clancy E, Kohli P, Jumper J, Hassabis D. 2022. Protein complex prediction with AlphaFold-Multimer. bioRxiv doi:10.1101/2021.10.04.463034:2021.10.04.463034.

37. Du Y, Hou G, Zhang H, Dou J, He J, Guo Y, Li L, Chen R, Wang Y, Deng R, Huang J, Jiang B, Xu M, Cheng J, Chen GQ, Zhao X, Yu J. 2018. SUMOylation of the m6A-RNA methyltransferase METTL3 modulates its function. Nucleic Acids Res 46:5195–5208.

38. Feederle R, Mehl-Lautscham AM, Bannert H, Delecluse HJ. 2009. The Epstein-Barr virus protein kinase BGLF4 and the exonuclease BGLF5 have opposite effects on the regulation of viral protein production. J Virol 83:10877–91.

39. Li R, Wang L, Liao G, Guzzo CM, Matunis MJ, Zhu H, Hayward SD. 2012. SUMO binding by the Epstein-Barr virus protein kinase BGLF4 is crucial for BGLF4 function. J Virol 86:5412–21.

40. Li R, Zhu J, Xie Z, Liao G, Liu J, Chen MR, Hu S, Woodard C, Lin J, Taverna SD, Desai P, Ambinder RF, Hayward GS, Qian J, Zhu H, Hayward SD. 2011. Conserved herpesvirus kinases target the DNA damage response pathway and TIP60 histone acetyltransferase to promote virus replication. Cell Host Microbe 10:390–400.

41. Li R, Liao G, Nirujogi RS, Pinto SM, Shaw PG, Huang TC, Wan J, Qian J, Gowda H, Wu X, Lv DW, Zhang K, Manda SS, Pandey A, Hayward SD. 2015. Phosphoproteomic Profiling Reveals Epstein-Barr Virus Protein Kinase Integration of DNA Damage Response and Mitotic Signaling. PLoS Pathog 11:e1005346.

42. Lv DW, Zhang K, Li R. 2018. Interferon regulatory factor 8 regulates caspase-1 expression to facilitate Epstein-Barr virus reactivation in response to B cell receptor stimulation and chemical induction. PLoS Pathog 14:e1006868.

43. Li J, Walsh A, Lam TT, Delecluse HJ, El-Guindy A. 2019. A single phosphoacceptor residue in BGLF3 is essential for transcription of Epstein-Barr virus late genes. PLoS Pathog 15:e1007980.

44. Zhang K, Lv DW, Li R. 2019. Conserved Herpesvirus Protein Kinases Target SAMHD1 to Facilitate Virus Replication. Cell Rep 28:449–459 e5.

45. Baquero-Perez B, Antanaviciute A, Yonchev ID, Carr IM, Wilson SA, Whitehouse A. 2019. The Tudor SND1 protein is an m(6)A RNA reader essential for replication of Kaposi’s sarcoma-associated herpesvirus. Elife 8.

46. Cianfrocco MA, Wong M, Youn C, Wagner R, Leschziner AE. 2017. COSMIC²: A Science Gateway for Cryo-Electron Microscopy Structure Determination. Practice & Experience in Advanced Research Computing New Orleans, LA doi:https://doi.org/10.1145/3093338.3093390.

47. Cianfrocco MA, Wong M, Youn C. 2017. COSMIC² – A science gateway for cryo-electron microscopy. Gateways 2017 Ann Arbor, MI

48. Pettersen EF, Goddard TD, Huang CC, Meng EC, Couch GS, Croll TI, Morris JH, Ferrin TE. 2021. UCSF ChimeraX: Structure visualization for researchers, educators, and developers. Protein Sci 30:70–82.

