## Supplemental Table 1 for "SUMOylation of the m6A reader YTHDF2 by PIAS1 promotes viral RNA decay to restrict EBV replication"

Table S1. Primers used in this study.

| Primer name | 5`-3` sequences |
| --- | --- |
| PSF010 (YTHDF2 K571R F) | cttgacgttcctttctaacactttcttcttcctcttggcg |
| pSF011 (YTHDF2 K571R R) | cgccaagaggaagaagaaagtgttagaaaggaacgtcaag |
| pSF029 (YTHDF2 K572R R) | gaccttgacgttcccttttaacactttcttcttcctcttggc |
| pSF030 (YTHDF2 K572R F) | gccaagaggaagaagaaagtgttaaaagggaacgtcaaggtc |
| pSF031 (YTHDF2 K281R F) | gcaacgggacccctgttatcccaagttccaatatcc |
| pSF032 (YTHDF2 K281R R) | ggatattggaacttgggataacaggggtcccgttgc |
| pSF048 (YTHDF2 KK-571/572-RR F) | accttgacgttcccttctaacactttcttcttcctcttggcg |
| pSF049 (YTHDF2 KK-571/572-RR R) | cgccaagaggaagaagaaagtgttagaagggaacgtcaaggt |
| pFS_38_F (pHTN-V5-YTHDF1 F insert) | atgggtaagcctatccctaaccctctcctcggtctcgattctacgatgtcggccaccagcgtggacacc |
| pFS_38_R (pHTN-V5-YTHDF1 R insert) | ggcccaaatctagatatccgtcattgtttgtttcgactct |
| pFS_39_F (pHTN-V5-YTHDF1 F vector) | agagtcgaaacaaacaatgacggatatctagatttgggcc |
| RL0325 (pHTN R vector) | cgtagaatcgagaccgaggagagggttagggataggcttacccatcggttgagctctgaattcggaagcgat |
| pFS_41_F (pHTN-V5-YTHDF3 F insert) | atgggtaagcctatccctaaccctctcctcggtctcgattctacgatgtcagccactagcgtggatcaga |
| pFS_41_R_N (pHTN-V5-YTHDF3 R insert) | ggcccaaatctagatatccgttattgtttgtttctatttctctccctacgcatggc |
| pFS_42_F_N (pHTN-V5-YTHDF3 F vector) | gccatgcgtagggagagaaatagaaacaaacaataacggatatctagatttgggcc |
| pFS_222_F (pLenti-YTHDF3 F) | gatctgccgccgcgatcgccatgtcagccactagcgtggatcaga |
| pFS_222_R (pLenti-YTHDF3 R) | tcgagcggccgcgtacgcgtttgtttgtttctatttctctccctacgcatggc |
| pFS_235_F (pLenti-YTHDF1 F) | gatctgccgccgcgatcgccatgtcggccaccagcgtggacacc |
| pFS_235_R (pLenti-YTHDF1 R) | tcgagcggccgcgtacgcgtttgtttgtttcgactct |
| pFS_232_F (YTHDF1 K227R F) | cacaggccccctgttatcccaggtgccaat |
| pFS_232_R (YTHDF1 K227R R) | attggcacctgggataacagggggcctgtg |
| pFS_233_F (YTHDF1 K551R F) | tctgccgttccctgcgcaccacctcc |
| pFS_233_R (YTHDF1 K551R R) | ggaggtggtgcgcagggaacggcaga |
| pFS_234_F (YTHDF3 K282R F) | ttaccactgaccctctttcatcccaagttccaatattcatgt |
| pFS_234_R (YTHDF3 K282R R) | acatgaatattggaacttgggatgaaagagggtcagtggtaa |
| pFS_237_F (18s RNA F): | ggttcgattccggagaggg |
| pFS_237_R (18s RNA R): | tcgggagtgggtaatttgc |
